# The *Vibrio cholerae* Type VI Secretion System Can Modulate Host Intestinal Mechanics to Displace Commensal Gut Bacteria

**DOI:** 10.1101/226472

**Authors:** Savannah L. Logan, Jacob Thomas, Jinyuan Yan, Ryan P. Baker, Drew S. Shields, Joao B. Xavier, Brian K. Hammer, Raghuveer Parthasarathy

## Abstract

Host-associated microbiota help defend against bacterial pathogens; the mechanisms that pathogens possess to overcome this defense, however, remain largely unknown. We developed a zebrafish model and used live imaging to directly study how the human pathogen *Vibrio cholerae* invades the intestine. The gut microbiota of fish mono-colonized by commensal strain *Aeromonas veronii* was displaced by *V. cholerae* expressing its Type VI Secretion System (T6SS), a syringe-like apparatus that deploys effector proteins into target cells. Surprisingly, displacement was independent of T6SS-mediated killing of *Aeromonas*, driven instead by T6SS-induced enhancement of zebrafish intestinal movements that led to expulsion of the resident commensal by the host. Deleting an actin crosslinking domain from the T6SS apparatus returned intestinal motility to normal and thwarted expulsion, without weakening *V. cholerae′s* ability to kill *Aeromonas in vitro*. Our finding that bacteria can manipulate host physiology to influence inter-microbial competition has implications for both pathogenesis and microbiome engineering.

## Introduction

The consortium of microbes that make up the human microbiome plays important roles in health and disease (1, 2). In the gastrointestinal tract, where most animal-associated microbiota reside and where the potential interface of interspecies contact is large, commensal microbes prevent colonization by pathogens, a function termed colonization resistance (3–6). Colonization resistance can, however, be thwarted by pathogens as the first stage of infectious disease, but the mechanisms used in this inter-species competition remain unclear. By understanding how pathogens interact with commensal communities, we may more rationally design future therapies focused on targeting the pathogens themselves, or on engineering the host microbiome to better resist disruption.

Uncovering these mechanisms, however, has proven challenging due to the difficulties of *in situ* monitoring of intestinal microbial populations and precise control of expression of pathogenic phenotypes.

We consider the transient human pathogen *Vibrio cholerae*, which can successfully colonize the human gut following ingestion of contaminated food or water. There, it causes diarrhea that may return the microbe to aquatic reservoirs in even larger numbers, leading to outbreaks. Cholera diarrhea causes severe dehydration and can be fatal if untreated. Recent epidemics in Haiti and Africa highlight that *V. cholerae* remains a major global problem and underscore that a better mechanistic understanding of the lifestyle of this microbe can help control future cholera outbreaks and infection (7).

*V. cholerae* can form biofilms on chitinous substrates such as the exoskeleton of crustaceans (8) and can colonize the gut of birds (9) and fish (10), which may promote transmission in aquatic environments. Within a human host, a complex set of signaling systems and external cues regulate colonization and disease factors, such as biofilm formation, chemotaxis-guided flagella, toxin-coregulated pili, several adhesins, and cell shape features, to ensure the microbe’s access to the intestinal surface (11, 12). Toxigenic isolates that carry the CTXphi prophage secrete the potent cholera toxin, which triggers rapid fluid loss and massive diarrhea. While cholera toxin itself serves as a competition factor by promoting dispersal of gut commensals, less is understood regarding additional factors that enable *V. cholerae* cells entering the gut to compete with the daunting assemblage of gut microbiota they encounter. Recent human studies show that cholera diarrhea disturbs the composition of the commensal microbiota (13) and studies in humans combined with mammalian animal models suggest that the microbiome composition affects how the host recovers from the disease (14).

Here, we sought to discover how *V. cholerae* may overcome resident commensals to invade a host intestine. We focused on the role of the type VI secretion system (T6SS), a syringe-like protein apparatus present in nearly 25% of all Gram-negative bacteria that inflicts damage on target cells by direct contact. The T6SS spike and inner tube pierce adjacent cells and deliver multiple “effector” molecules that can be deadly to eukaryotic cells, as well as bacteria that lack immunity proteins (15–17). T6 activity in non-toxigenic, environmental isolates and in toxigenic, CTXphi isolates derived from clinical sources is controlled by diverse regulatory systems and external cues (18, 19). Recently, a role for T6-mediated microbe-microbe interactions within the mouse gut has been demonstrated for *Shigella* and *Salmonella* infection (20, 21). Commensal *Bacteroides* can use their T6SS to compete with other bacteria to maintain their presence in the mouse gut (22, 23). T6SS genes have been detected in the human gut microbiome as well (24, 25). All of this evidence suggests that T6SSs require more attention for their role in the initiation and development of cholera, and also in mediating microbe-microbe and microbe-host interactions in the gut microbiome.

Investigating the potential role of the *V. cholerae* T6SS in intestinal invasion is challenging in humans, and even in mammalian model organisms, due to the complexity of colonization and infection processes and the severe difficulty of *in vivo* imaging. By contrast, zebrafish are a powerful laboratory model for the direct observation and experimental control of microbiome interactions. Germ-free and gnotobiotic protocols allow precise control of intestinal microbial composition (26). Zebrafish are relatively transparent at larval stages. Thus, light sheet fluorescence microscopy (27–29) can be used to capture detailed three-dimensional images of fluorescently-labeled bacteria, spanning the entire gut, over many hours, to monitor both sudden and longer-term transitions in bacterial populations (30), and differential interference contrast microscopy can capture the dynamics of unlabeled intestinal tissue in the same animal (31).

Mammalian models for *V. cholerae* infection have revealed modest contributions of the T6SS in the infant rabbit (32), and fluid accumulation in the infant mouse (33). However, these organisms, unlike fish (34) or humans, are not natural *V. cholerae* hosts (35). Recent studies have demonstrated the utility of the zebrafish as a model for cholera intestinal colonization, pathogenesis, and transmission (36), revealing for example that fish colonization is independent of cholera toxin (37). Together, these features make the zebrafish an ideal model for studying the dynamics of vertebrate gut colonization by *Vibrio cholerae*, and specifically the role of its T6SS.

In this study, we combined microbial genetics, *in vitro* experiments and quantitative *in vivo* imaging in zebrafish to determine the role of the T6SS of *V. cholerae* in gut colonization. We exploited the known regulation pathways of T6SS (38–40) to genetically manipulate the human-pathogenic *V. cholerae* wild type El Tor strain C6706 to constitutively express functional, defective, or altered T6SS machinery, as well as generating strains lacking T6SS immunity. We then imaged at high resolution the invasion by *V. cholerae* of zebrafish intestines that were previously colonized by a zebrafish-commensal *Aeromonas* species. Our experiments show a strongly T6SS-dependent displacement of the resident bacteria. The displacement took the form of sudden collapses in *Aeromonas* populations via ejections of aggregated bacteria from the gut, similar to the collapses previously reported for *Aeromonas* when challenged by a fish-commensal species of the genus *Vibrio* (30). We found that the expression by *V. cholerae* of a functional T6SS induced a large increase in the amplitude of the peristaltic movements in the host intestine. Deletion of the actin cross-linking domain (ACD) of one of the T6SS spike proteins returned zebrafish gut activity to normal and eliminated *V. cholerae′s* ability to expel the commensal *Aeromonas* from the gut, without affecting its ability to kill *Aeromonas* cells *in vitro*.

To the best of our knowledge, ours is the first observation that the bacterial T6SS can induce organ-level physiological changes in an animal host that displace resident microbiota. These findings expand the array of known molecular mechanisms by which pathogens can leverage host-microbe interactions to redefine microbial community composition, and also suggest that the T6SS could be rationally manipulated to deliberately engineer the human microbiome.

## Results

### Human-derived *Vibrio cholerae* colonizes the larval zebrafish intestine but exhibits weak intra-species T6SS-mediated killing *in vivo*

A streptomycin resistant mutant of patient-derived El Tor biotype C6706 served as a “wild type” strain (denoted T6SS^WT^) as it is proficient at T6-mediated bacterial killing (39, 41). T6SS and immunity genes are well characterized in this strain, allowing us to construct variants that differed in T6SS expression, immunity, and functionality (Fig. 1A). A strain constitutive for T6SS expression, termed T6SS^+^, was previously constructed by replacement of the native *qstR* promoter, and a T6SS^−^ derivative of this strain was constructed by deletion of the *vasK* gene (Δ*vasK*) (42). Further deletion of three T6 immunity genes (*tsiV1-3*) generated a T6SS^−^ Imm^−^ strain. Each strain was labeled fluorescently either with a chromosomally-introduced teal or orange fluorescent protein to enable microscopy (42). (Supp. Table).

**Figure 1.**
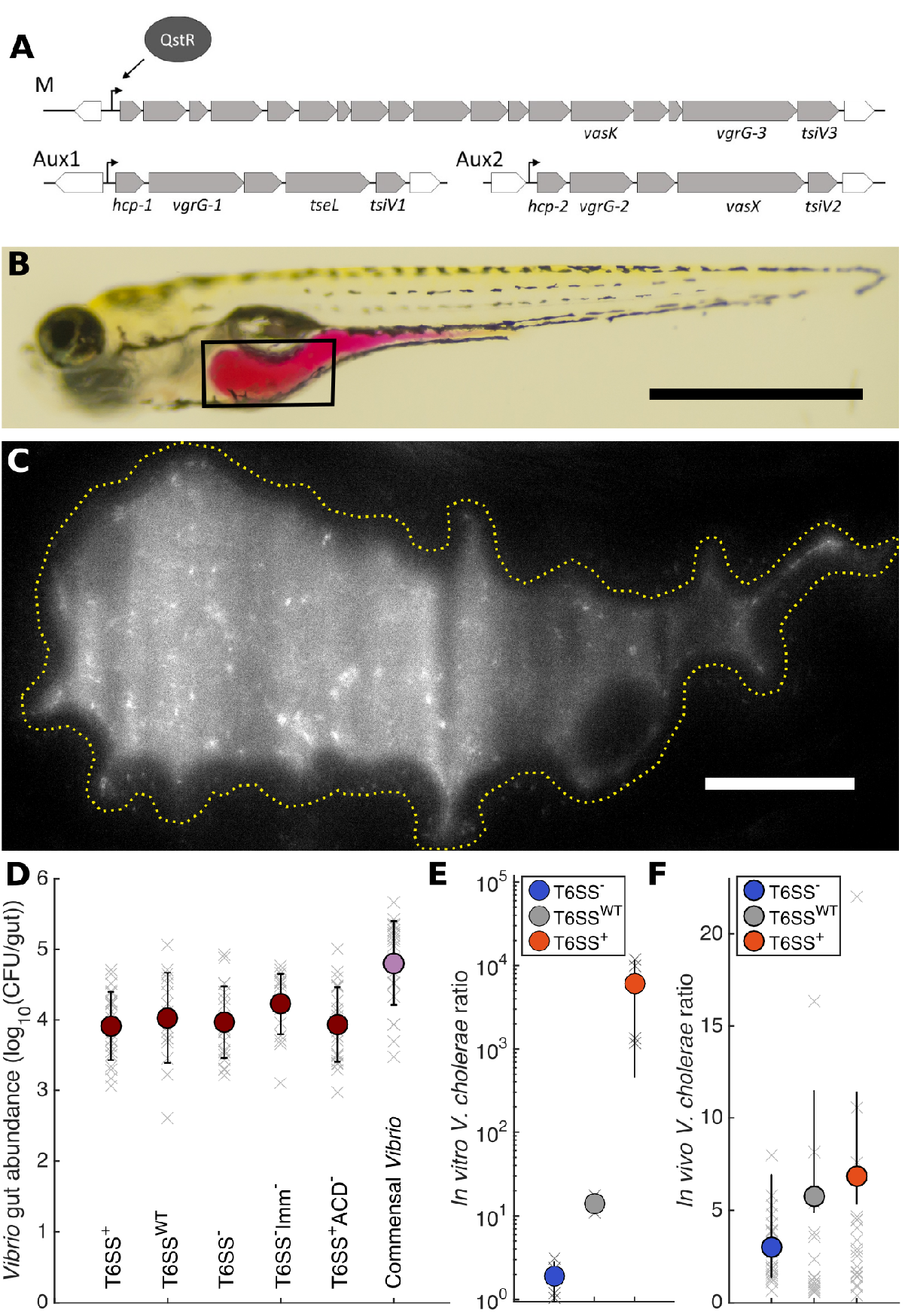
(A) Genes of the *V. cholerae* C6706 Type VI secretion system. T6SS genes are primarily organized in three operons that are transcriptionally activated through the regulator QstR. The main cluster (M) encodes most of the T6SS structural genes while the major Hcp subunit is encoded on the auxiliary clusters Aux1 and Aux2. Each cluster terminates in genes encoding antibacterial effectors (TseL, VasX and VgrG-3) and their respective immunity proteins (TsiV1-3). Each cluster also encodes proteins that form a spike at the apex of the apparatus, VgrG-1-3. Two of the three VgrG proteins are multifunctional: VgrG1 contains a C-terminal actin crosslinking domain (ACD), and VgrG-3 has a muramidase domain that serves as an antibacterial effector. (B) A larval zebrafish at 5 dpf with the intestine colored for illustration by orally gavaged phenol red dye. Scale bar: 1mm. (C) A light sheet fluorescence image of wild type *V. cholerae* expressing orange fluorescent protein in the larval zebrafish intestine. The region shown roughly corresponds to the box in (B), with the luminary boundary roughly indicated by the yellow dotted line. Individual motile bacteria are evident, as is the background autofluorescence of the gut lumen. See also Supplemental Movie 1. Scale bar: 50 μm. (D) Abundance of *Vibrio* strains in the larval zebrafish intestine at 24 hours post-inoculation. All *V. cholerae* strains robustly colonize to approximately 10^4^ bacteria per fish, roughly an order of magnitude lower than a commensal *Vibrio* species (rightmost data points). Measurements from individual fish at 6 dpf are shown in grey, averages are indicated by solid colored circles, and black error bars represent quartiles. (E) Ratios of *V. cholerae* strains in an *in vitro* competition assay. Each indicated strain was mixed 1:1 with T6SS^−^ Imm^−^ as a target, and spotted onto a nylon membrane on agar. Ratios were determined from CFU counts following 3h of incubation. The T6SS+ strain exhibits a large competitive advantage over the T6SS^−^ Imm^−^ strain. (F) Ratios of *V. cholerae* strains in the larval zebrafish intestine 24 hours after co-inoculation. At 5 dpf, fish were coinoculated with T6SS^−^ Imm^−^ as a target, and one of either wild type, T6SS^−^, or T6SS+ strains. The T6SS+ and wild type strains exhibit a slightly greater competitive advantage over the T6SS^−^ Imm^−^ strain compared to the T6SS^−^ strain. In (E) and (F), measurements from individual fish are shown in grey, averages are indicated by solid circles, and quartiles are represented by black lines.

To determine whether the human-derived *V. cholerae* and its variants could colonize the larval zebrafish gut, we inoculated flasks housing germ-free larvae with a single bacterial strain at 5 days post-fertilization (dpf). We then dissected the gut at 6 dpf and determined intestinal bacterial abundance by serial plating and counting colony forming units (CFUs). For comparison, we also considered a previously examined zebrafish commensal bacterium ZWU0020 assigned to the genus *Vibrio* (30, 43). All *V. cholerae* strains examined could colonize the larval zebrafish intestine robustly to an abundance of approximately 10^4^ CFU per gut, which is roughly one order of magnitude lower than the commensal *Vibrio* (Fig. 1B-D). Direct observation by light sheet fluorescence microscopy at 5 dpf showed that each strain of *V. cholerae* was abundant and highly motile in the intestinal lumen (Fig. 1C and Supplemental Movie 1).

We then asked whether we could detect signatures of T6SS-mediated intra-species competition *in vitro* and *in vivo*. For *in vitro* assays, we mixed two *V. cholerae* strains in liquid culture at a 1:1 ratio. One of these was always the T6SS^−^Imm^−^ strain which, lacking immunity to T6SS, served as a “target” for inter-bacterial killing (39). We spotted the mix onto nylon membranes on agar plates, and allowed the microbes to interact in close proximity for 3 hours. We then quantified killing by measuring ratio of CFU counts for each pair of strains, which we distinguished by their fluorescent markers. T6SS^−^ and T6SS^WT^ strains were only slightly enhanced compared to the target; the T6SS^+^ strain, however, dominated the mixture, indicating T6SS-mediated killing (Fig. 1E), consistent with prior in vitro work (39). *In vivo*, we co-inoculated zebrafish flasks at 5 dpf with the orange-labeled T6SS^−^Imm^−^ strain and one of either the teal-labeled wild type, T6SS^−^ defective, or T6SS+ constitutive strains at a 1:1 initial ratio. We determined their ratios in the fish at 6 dpf using gut dissection and plating, again differentiating the strains by their fluorescence (Fig. 1F). We found that the T6SS+ and wild type strains, compared to the T6SS^−^ strain, exhibited a small and variable competitive advantage over the T6SS^−^Imm^−^ target strain, with abundance ratios of 6.8 ± 2.9, 5.8 ± 2.6, and 3.0 ± 0.4 for challenge by T6SS+, T6SS^WT^, and T6SS^−^, respectively (mean ± s.e.m.). The *in vivo* killing rate by T6SS competent cells was only roughly a factor of 2 higher for strains with functional T6SS compared to strains without the functional T6SS (Fig. 1F); this is far less dramatic than the *in vitro* killing of *V. cholerae* by other *V. cholerae* (Fig. 1E).

### Constitutive expression of T6SS potentiates *Vibrio cholerae* invasion of zebrafish intestines occupied by a commensal species

Next, we addressed the key question of whether the T6SS can affect the ability to invade an established, commensal intestinal microbial community. We used as our target species *Aeromonas veronii* strain ZOR0001, hereafter referred to as *Aeromonas*, a Gram-negative bacterium native to and commonly found in the zebrafish intestine (44). Prior work has shown that this strain can robustly mono-colonize germ-free larval zebrafish at 10^3^-10^5^ bacteria per gut (29, 43). *Aeromonas* forms dense bacterial aggregates *in vivo* (29), and can be invaded by the fish-commensal *Vibrio* sp. ZWU0020 (30).

We first determined whether *Aeromonas* was susceptible to T6-mediated killing by *V. cholerae in vitro*. We mixed *Aeromonas* and *V. cholerae* strains in liquid culture and spotted them onto nylon membranes as in the previously described *in vitro* experiments. We then quantified killing by measuring *Aeromonas* CFU counts before and after the membrane incubation. Aeromonas CFU counts when mixed with T6SS^−^ *V. cholerae* were indistinguishable from those of a control mix of *Aeromonas* with *Aeromonas* (Fig. 2A). Wild type *V. cholerae*, and particularly the T6SS+ strain, decreased Aeromonas CFU counts significantly, indicating high inter-species killing rates (Fig. 2A, Supp. Movies 2-4), consistent with prior *in vitro* results with an *Escherichia coli* target (19).

**Figure 2.**
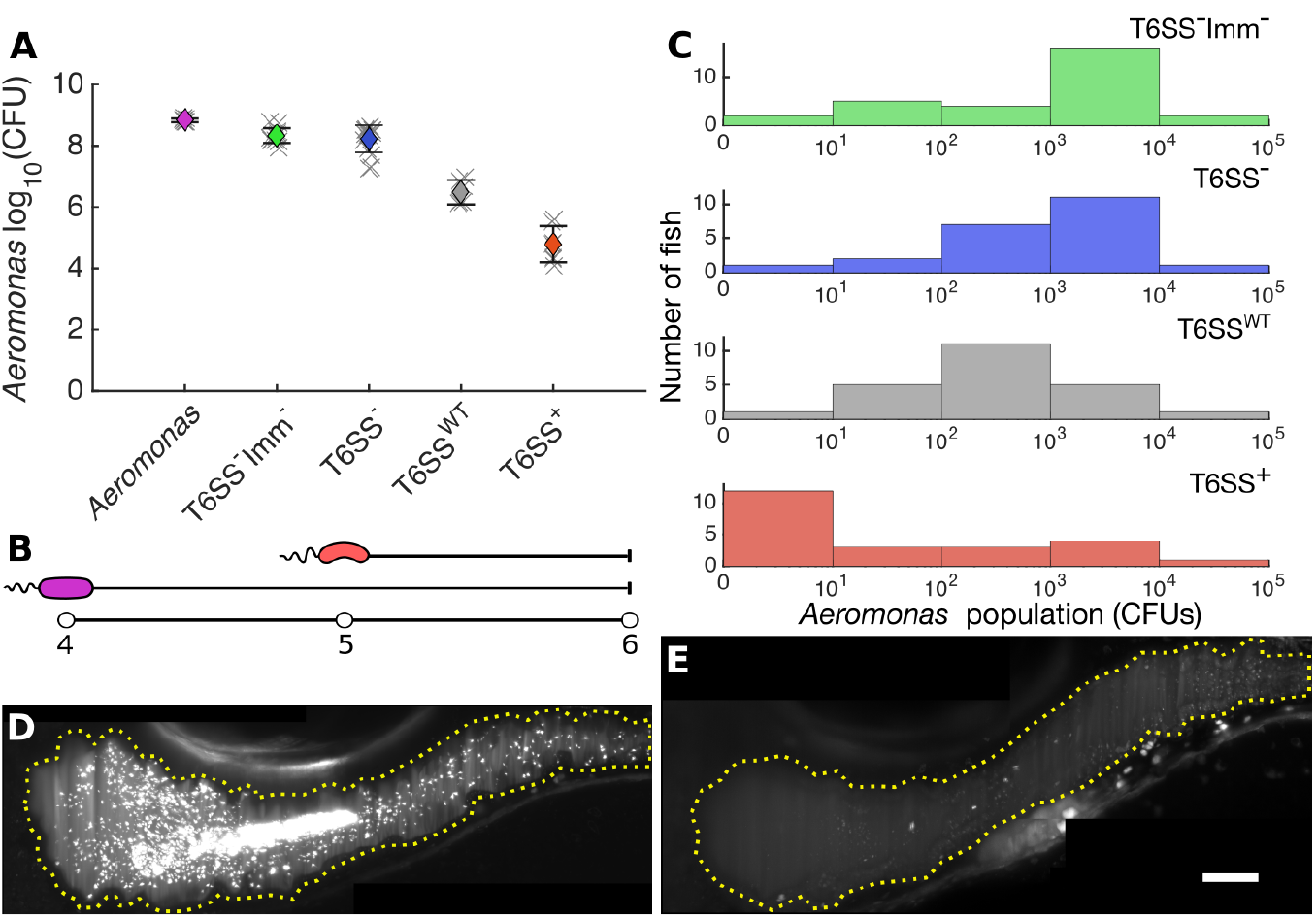
(A) *In vitro* abundances of *Aeromonas* when mixed with *Vibrio cholerae* strains, determined from spotting liquid-cultured pairs of strains onto agar-supported membranes, 3h after mixing. (B) Schematic diagram of the protocol used to characterize *Aeromonas—Vibrio* interactions *in vivo. Aeromonas* (magenta) is allowed to colonize at 4 dpf followed by inoculation of *V. cholerae* (orange) strains into the surrounding water at 5 dpf. Imaging and/or dissections and serial plating occur at 6 dpf. (C) Histogram of *Aeromonas* abundances in the larval gut 24 hours after potential invasion by *V. cholerae* strains. The peak abundances are roughly 10^3^ CFU/gut when *Aeromonas* is followed by T6SS^−^ Imm^−^ and T6SS, 10^2^ when followed by wild type, and 0 when followed by T6SS+. (D, E) Maximum intensity projections of a 3D light sheet image stack of *Aeromonas* in the larval gut 24 hours after invasion by T6SS^−^ Imm^−^ (D) and T6SS+ *V. cholerae* (E) with the boundaries of the gut lumen roughly indicated by yellow dotted lines. Scale bar: 50 μm.

To determine the role of the T6SS *in vivo*, we monocolonized zebrafish by inoculating flasks containing germ-free larvae with *Aeromonas* at 4 dpf, and then inoculated with one of the *V. cholerae* strains at 5 dpf (Fig. 2B, N~30 animals per *V. cholerae* strain). Gut dissection and serial plating at 6 dpf revealed dramatic differences in the *Aeromonas* abundance depending on the T6SS of the invading strain. *Aeromonas* challenged by T6SS^−^ or T6SS^−^ Imm^−^ *V. cholerae* persisted in the gut at approximately 1000 CFU per fish on average (Fig. 2C, first and second panels). *Aeromonas* challenged by the T6SS+ *V. cholerae*, however, fell to single-digit numbers, with zero detectable *Aeromonas* in over 50% of fish (Fig. 2C, bottom panel). *Aeromonas* challenged by the wild type *V. cholerae* showed intermediate numbers between the T6SS^−^-challenged and the T6SS+-challenged strains (Fig. 2C, third panel). Live imaging 24 hours after the *V. cholerae* inoculation demonstrates the differential impacts on the resident *Aeromonas*, with large populations consisting of dense clusters and discrete individuals in the gut of larvae challenged by T6SS^−^ *V. cholerae* (Fig. 2D), and few *Aeromonas* remaining in the gut of larvae challenged by T6SS^+^ *V. cholerae* (Fig. 2E). Each of the invading *V. cholerae* strains was present at 6 dpf at approximately 10^4^ CFU/gut (Suppl. Fig. 1B).

### *Aeromonas* are expelled in frequent sudden collapses from fish guts invaded by T6SS-expressing *V. cholerae*

To better characterize the strong T6SS-mediated effect of *V. cholerae* on gut-resident *Aeromonas*, we monitored bacterial population dynamics over 12-17 hour durations using light sheet fluorescence microscopy, capturing a three-dimensional image spanning the entire larval intestine every 20 minutes. We used the same inoculation protocol and began live imaging 8 hours after *V. cholerae* inoculation.

We had shown in previous work that *Aeromonas* populations residing the zebrafish intestine can be punctuated by occasional large collapses corresponding to ejection from the gut. In fish monocolonized with *Aeromonas*, these collapses occurred at a mean rate of *p*_c_ = 0.04 ± 0.02 hr ^−1^, but in fish invaded by the commensal *Vibrio* ZWU0020 the rate of collapse increased to *p*_c_ = 0.07 ± 0.02 hr^−1^ (30). Here, as in prior work, we defined a collapse as a population drop of at least a factor of ten in one 20-minute time interval, together with at least a factor of two drop relative to the original population at the subsequent time step. The *Aeromonas* population was strikingly stable when invaded by the T6SS^−^ strains: we observed zero collapses during the entire 58.0 and 70.3 hour total imaging times for T6SS^−^ Defective (*N* = 5 fish) and T6SS^−^ Immunity-(*N* = 6 fish) *V. cholerae* challenges, respectively (Fig. 3A, first two panels). Challenge by the wild type *V. cholerae* resulted in 2 population collapses in 72.7 hours corresponding to a collapse rate *p*_c_ = 0.03 ± 0.02 hr ^−1^ (Fig. 3A, third panel, *N* = 6 fish). Challenge by T6SS+ *V. cholerae* gave rise to large and frequent collapses, totaling 8 in 64.3 hours (Fig. 3A, last panel, *N* = 5 fish), corresponding to a collapse rate *p*_c_ = 0.12 ± 0.04 hr ^−1^, nearly twice as large as that induced by the fish-commensal *Vibrio* ZWU0020 (30) (Fig. 3B-D; Supplemental Movies 5-6).

**Figure 3.**
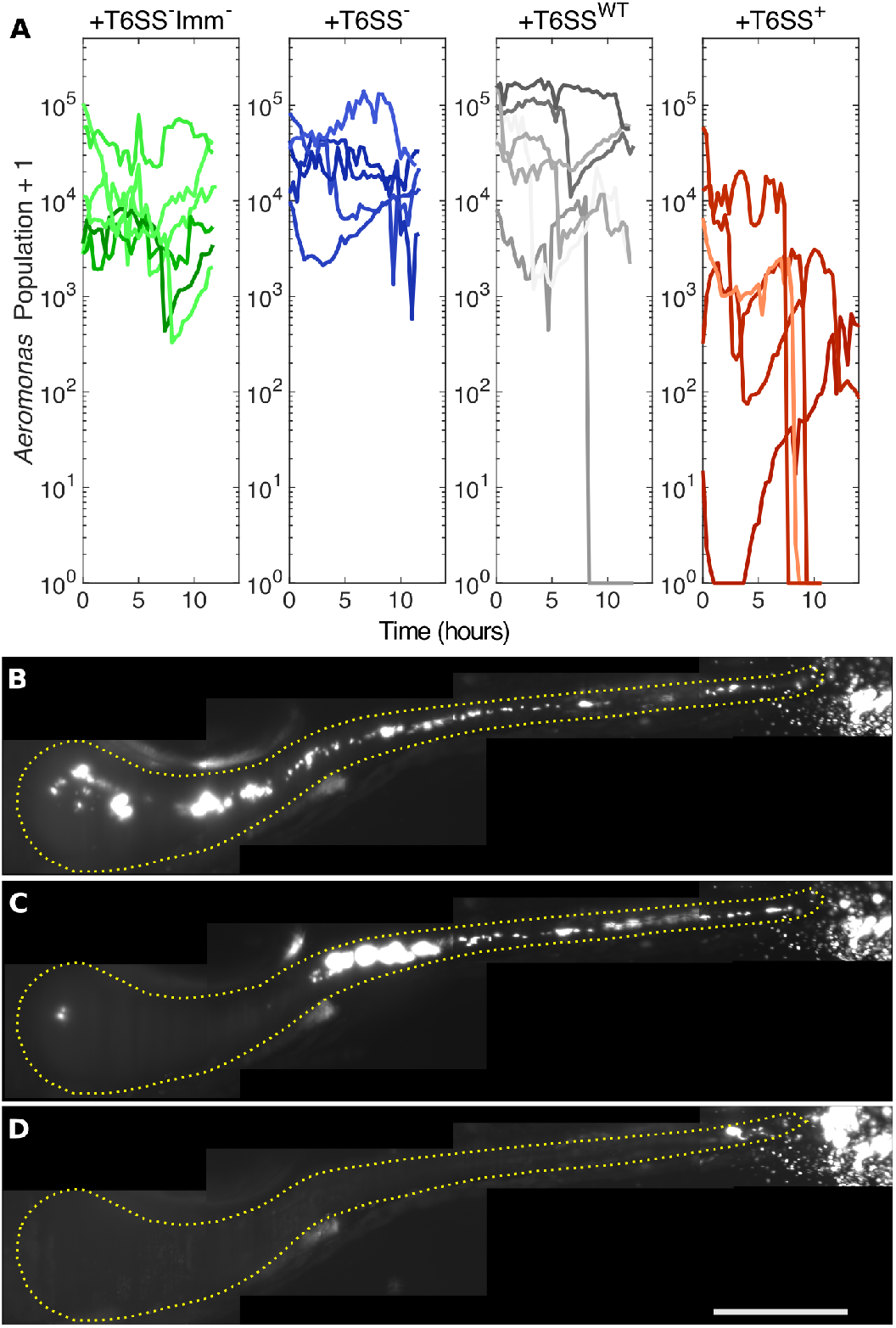
(A) Time-series of *Aeromonas* populations in larval zebrafish intestines when challenged by different strains of *Vibrio cholerae*, derived from light sheet fluorescence imaging. Beginning 8 hours after *Vibrio* inoculation, fish were imaged every 20 minutes for 12-17 hours. Each curve is from a different zebrafish. When invaded by T6SS+ *V. cholerae*, overall *Aeromonas* abundance is low, and collapses in population of over an order of magnitude are evident. (B,C,D) Maximum intensity projections of a 3D light sheet image stack of *Aeromonas* in a larval zebrafish intestine invaded by T6SS+ *V. cholerae* at 9.3, 10.7, and 16.3 hours after the start of imaging. A collapse of the *Aeromonas* population is evident as time progresses. See also Supplemental Movies 5-6. Yellow dotted lines roughly indicate luminary boundary. Scale bar: 200 μm.

### Constitutively expressed T6SS alters the intestinal movements of larval zebrafish in an ACD-dependent manner

The larval zebrafish intestine, like those of other animals, has periodic propagative contractions that drive the motion of dense aggregates of *Aeromonas* and can ultimately cause their ejection (30). We tested whether the collapses in the *Aeromonas* populations observed in the T6SS+ competition (Fig. 3B-D) could be due to greater gut motility. We compared intestinal movements of germ-free fish and fish mono-associated with the various *V. cholerae* strains using differential interference contrast microscopy (DICM), which allowed direct visualization of the intestinal epithelial tissue and lumenal space (Fig. 4A) (31). Then, we used image velocimetry techniques to quantify the the frequency and amplitude of intestinal contractile waves (Fig. 4B-D) (30, 45). None of the strains altered the frequency of peristaltic contractions, compared to germ-free fish (Fig. 4E). The amplitude of the contractions, however, was greatly enhanced in the fish colonized with T6SS^+^ strain, but not other strains (Fig. 4F, Supplemental Movies 7-8). The magnitude of the effect, roughly a 200% increase in the amplitude of contractions compared to germ-free fish, was remarkable and unexpected. For comparison, treatment with the neurotransmitter acetylcholine or deletion of all enteric neurons induces at most a change of roughly 40% in peristaltic amplitude (45).

Though this T6SS-dependent alteration of host gut motility was unexpected, there are well-established precedents for T6SS-mediated *V. cholerae* interactions with eukaryotic cells driven by an actin crosslinking domain (ACD) present in the C-terminus of the VgrG-1 spike protein of the T6 secretion apparatus (33). We hypothesized that the ACD might also be responsible for the larger amplitude of gut motility. To test this hypothesis, we deleted the ACD of *vgrG-1* in the constitutive T6SS-expressing *V. cholerae* T6SS+ strain (see Methods). When we mono-colonized zebrafish larvae with T6SS^+^ ACD^−^ *V. cholerae*, we observed no increase in either the frequency or amplitude of intestinal contractions compared to germ-free fish (Fig. 4E,F). This strain, however, maintained the ability to kill *Aeromonas in vitro* at a rate similar to that of T6SS^+^ strain, which indicates an otherwise functional T6SS (Fig. 4G). Therefore, the VgrG-1 ACD is specifically necessary for the increase of amplitude in intestinal contractions observed in fish mono-colonized with the T6SS^+^ strain.

**Figure 4.**
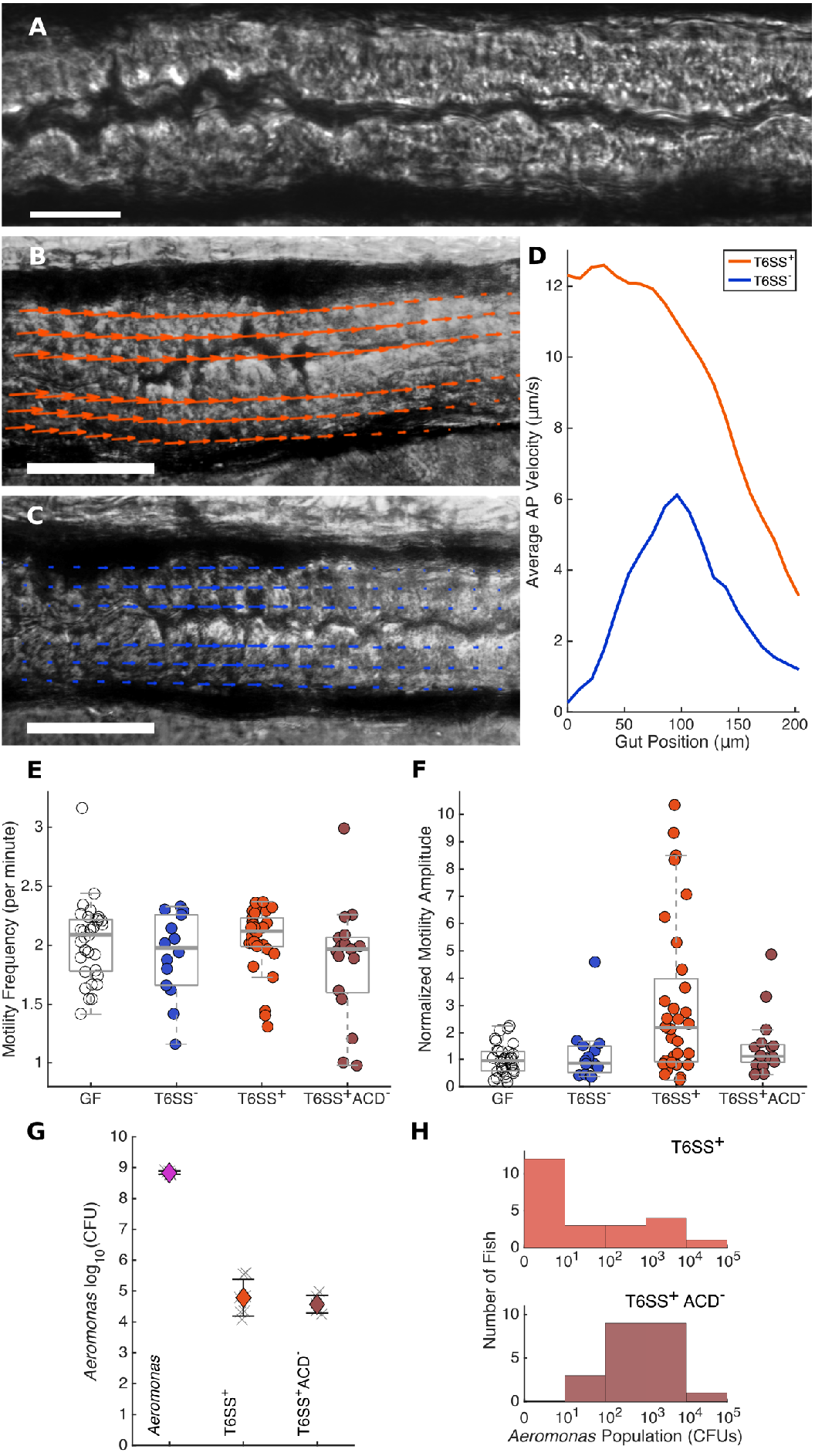
(A) A DIC image of a portion of a larval zebrafish intestine. Scale bar: 50 μm. See also Supplemental Movies 7-8. (B, C) DIC images of portions of larval zebrafish intestines with superimposed image velocimetry vectors indicating intestinal motions, from fish mono-associated with T6SS+ (B) and T6SS^−^ (C) *Vibrio cholerae* strains. Scale bars: 50 μm. (D) Examples of average anterior-posterior velocities along the gut axis, corresponding to the fish in (B,C). (E) Frequency of periodic gut motility for germ-free fish and fish mono-associated with T6SS^−^, T6SS^+^, and T6SS^+^ACD^−^ strains. (F) Gut motility amplitudes under the same conditions as panel (B), normalized by the mean value in germ-free fish. Fish associated with T6SS^+^ show far greater gut motility amplitude than T6SS^−^, T6SS^−^ACD^−^, or germ-free fish. (G) *In vitro* killing rates of *Aeromonas* by T6SS^+^ and T6SS^+^ACD^−^ *Vibrio cholerae* strains. (H) Histogram of *Aeromonas* abundances in the zebrafish gut 24 hours after potential invasion by T6SS^+^ (the same data as in Figure 2C) and T6SS^+^ACD^−^ strains. The peak abundances are roughly 0 when followed by the T6SS^+^ strain, but 10^2^-10^3^ CFU/gut when followed by the T6SS^+^ACD^−^ strain.

We then tested the ability of T6SS^+^ ACD^−^ *V. cholerae* to invade an intestinal *Aeromonas* population using the same zebrafish invasion assay described above. While the *Aeromonas* population drops precipitously following T6SS^+^ *V. cholerae* introduction, *Aeromonas* after T6SS^+^ ACD^−^ *V. cholerae* introduction remained abundant, averaging approximately 1000 CFU per fish (N = 31 fish) similar to the numbers seen when challenged by T6SS^−^ strains (Fig. 4H). T6SS^+^ ACD^−^ *V. cholerae* was nonetheless present in the gut at high abundance, approximately 10^4^ CFU/gut (Supp. Fig. 1B). This experiment demonstrated that removing the T6SS actin crosslinking domain eliminates *V. cholerae’s* ability to displace a competitor, despite an otherwise functional T6SS capable of killing *in vitro*. Taken together, these results show that the ability of *V. cholerae* to dominate a gut colonized by *Aeromonas* works specifically by increasing the amplitude of host peristalsis in a manner dependent on the VgrG-1 ACD.

## Discussion

We have shown that *V. cholerae* can employ its type VI secretion system to amplify the intestinal contractions in zebrafish and induce the host to expel from its gut a resident microbiota of the commensal genus *Aeromonas*. The coupling of T6SS activity to host contractions depended on an actin crosslinking domain (ACD) of the T6SS apparatus; when the ACD was deleted, *V. cholerae* could no longer induce enhanced host contractions and dense *Aeromonas* communities remained in the gut. The effect on the host peristalsis was specific; deleting the ACD did not affect the ability of *V. cholerae* to kill *Aeromonas* on contact, nor did it impact the ability of *V. cholerae* to enter and occupy the host intestine. *V. cholerae* itself seemed unaffected by the enhanced intestinal motility, likely due to its ability to remain planktonic and motile inside the zebrafish gut (Fig. 1C, Supp. Movie 1), which is similar to what was seen previously for the highly motile commensal *Vibrio* ZWU0020 (30, 46). Taken together, our results show that an enteric colonizer (*V. cholerae*) can use a previously undiscovered host-microbe interaction (T6SS-dependent enhancement of gut contractions) to influence the population dynamics of a competitor (*Aeromonas*).

The existence of this inter-species interaction raises intriguing questions and opens avenues for future research. Perhaps at the forefront is the question of what leads T6SS-affected host cells, likely lining the gut, to unleash large contractions of the entire organ. The host cellular mechanism is likely complex and could involve one or more cell types: immune cells, which at larval stages in zebrafish include neutrophils and macrophages that can respond to microbial presence (43) and may take up T6SS-active intestinal bacteria; epithelial cells, a variety of which line the gut and lie in close proximity to lumenal bacteria; cells of the enteric nervous system, which include enteric neurons that coordinate muscle activity; and the smooth muscle cells that are the proximate cause of contractions. Identifying the cells and signaling mechanisms that link the bacterial T6SS to intestinal mechanics is a challenging but attainable goal for research groups in years to come. The tools of transgenic and gnotobiotic zebrafish together with three-dimensional live imaging and image analysis empower future studies to discover new links between cellular processes and macroscopic organ behavior.

This newly discovered T6SS function is important in its own right, standing out from previously identified mechanisms for orchestrating the ecology of the microbiome (47) because it highlights the role of host fluid-mechanical environment in shaping gut population dynamics, an emergent theme in contemporary microbiome research (30, 46, 48, 49). The T6SS itself has received well deserved attention for its dramatic role in contact-mediated interbacterial toxicity (50), and potential implications in mediating interbacterial competition within the animal microbiome (22, 24, 25). Given the prevalence of T6SS among bacteria, the possibility that gut-colonizing bacteria can use the T6SS to manipulate host peristalsis suggests that this could be a common tactic for bacteria to indirectly influence interbacterial competition. Moreover, exogenous delivery of T6SS proteins, or their engineering into otherwise beneficial microbes, could offer a new path to therapeutic modulation of human gastrointestinal activity.

Our observations may also inform our understanding of T6SS regulation. Previous studies have shown that chitinous material triggers T6SS activity in *V. cholerae* (19, 35, 39); chitin can be found in crab shells, zooplankton exoskeletons, and marine snow commonly colonized by *Vibrios* in aquatic environments (51, 52). We found that wild type C6706 *V. cholerae*, but not T6SS^−^ derivatives, was capable of modest killing of *Aeromonas in vitro* (Fig. 2A), consistent with results observed prior for C6706 T6 killing of an *E. coli* target (19). We also observed small reductions in *Aeromonas* counts and rare extinction events *in vivo* (Fig 2C and Fig. 3A). Since the germ-free zebrafish used here were not provided with a chitin source, it is interesting to speculate that endogenous chitin production recently documented within the juvenile zebrafish gut (53) is inducing the wild type *V. cholerae* T6-mediated activity observed here. Further studies will determine the contribution that chitin signaling plays in T6 expression by *V. cholerae* in fish intestinal environments.

Most directly, our work sheds a new light on the role of the T6SS in the ecological dynamics of *V. cholerae* colonization of a vertebrate host. The ability of the T6SS to amplify host intestinal mechanics was previously undetected, likely for three reasons. First, the development of cholera in humans is a complex, multifactorial process in which the role of T6SS may be confounded by other factors, most importantly the strong effects of the cholera toxin. Second, the animal models typically used in cholera research are not native cholera hosts, and the mechanisms of their colonization may be different. Fish, however, naturally host *V. cholerae*, and because zebrafish colonization depends less on other factors we could detect the effects of T6SS on intestinal movements, and we could then use genetically modified *V. cholerae* strains to confirm the molecular mechanism. Third, the zebrafish model allows direct, quantitative, *in vivo* imaging using modern microscopy methods, in contrast to indirect, static, DNA-or RNA-sequencing-based assays typically used to study mouse or human microbiomes. *In vivo* imaging greatly facilitates observations of intestinal activity and enabled the discovery of sudden spatiotemporal changes in bacterial distributions. How our findings may map onto *V. cholerae* colonization in humans is unknown, but a role for T6SS-mediated activity is certainly plausible. Establishing this will take further investigation, especially for the potential aim of designing therapeutics that target the T6SS to prevent colonization in humans or in environmental reservoirs such as fish. Nonetheless, our results enhance our understanding of the strategies and abilities of *V. cholerae*, a pathogen that continues to impact millions of people worldwide.

## Materials and Methods

### Ethics Statement

All experiments involving zebrafish were carried out in accordance with protocols approved by the University of Oregon Institutional Animal Care and Use Committee and followed standard methods (54).

### Gnotobiotic Techniques

Wild-type larval zebrafish were derived devoid of microbes as previously described in (26). In brief, fertile eggs were collected and placed in a sterile antibiotic embryo media solution consisting of 100 μg/ml ampicillin, 250 ng/ml amphotericin B, 10 μg/ml gentamycin, 1 μg/ml tetracycline, and 1 μg/ml chloramphenicol for approximately six hours. The eggs were then washed in a sodium hypochlorite solution and a polyvinylpyrrolidone-iodine solution. Washed embryos were distributed in sets of 15 into tissue culture flasks containing 15μl of sterile embryo media. Flasks of larval zebrafish were inspected for sterility prior to their use in experiments.

### Bacterial Strains and culture conditions

*Aeromonas veronii* (ZOR0001, PRJNA205571) and *Vibrio* (ZWU0020, PRJNA205585) were isolated from the zebrafish intestinal tract as previously described (44). These strains were fluorescently labeled with EGFP or dTomato for imaging experiments with methods similar to (55) as described in (30).

All *V. cholerae* strains were derivatives of El Tor C6706 str-2 (see Supplementary Table). Bacterial cultures were routinely grown at 30°C or 37°C in lysogeny broth (LB) with shaking, or statically on LB agar. In-frame deletion mutants and promoter-replacements in *V. cholerae* were constructed using the allelic exchange method described previously (Skorupski 1996). Standard molecular biology-based methods were utilized for DNA manipulations. DNA modifying enzymes and restriction nucleases (Promega and New England Biolabs), Gibson assembly mix (New England Biolabs), Q5, Phusion and OneTaq DNA Polymerases (New England Biolabs) were used following the manufacturer’s instructions. All recombinant DNA constructs were verified by Sanger sequencing (Eurofins).

### Culture-Based Quantification of Bacterial Populations

Germ-free larval zebrafish were inoculated with select bacterial strains as in previous work (30, 43). Bacteria were grown on a shaker in Luria Broth for 10-14 hours at 30 ^o^C. Bacteria were prepared for inoculation by pelleting for two minutes at 7000g and were washed once in sterile embryo media prior to inoculation. An inoculum of 10^6^ CFU/ml was used for *Aeromonas* (ZOR0001, PRJNA205571) and *Vibrio* (ZWU0020, PRJNA205585) strains and 10^7^ CFU/ml for *Vibrio cholerae* strains. Bacterial inoculums were added directly to tissue culture flasks containing germ-free larval zebrafish.

### In vitro measurements of bacterial competition

For *in vitro* killing assays, bacterial strains were inoculated from glycerol stock and shaken in lysogeny broth (LB) at 30 ^o^C or 37 ^o^C overnight. The cells were then harvested, washed in sterile phosphate buffered saline (PBS) twice and normalized to OD600=1 in PBS. Pairs of strains were mixed 1:1, and 25 μl of the liquid was spotted onto a 0.20 μm diameter porous nylon membrane filter (Millipore) placed on an LB agar plate. After allowing them to dry, plates were incubated at 37 ^o^C for 3h. Each membrane was then carefully removed from the agar plate and vortexed in sterile PBS for 1 min. The killing rate was assessed by comparing the target cell numbers before and after incubation by plating and counting colony forming units (CFUs). An antibiotic resistant marker (streptomycin or gentamicin) inserted into the target strain chromosome enabled discrimination of target cells for CFU counting.

For *in vitro* time lapse fluorescence microscopy, bacterial strains were inoculated from glycerol stocks and shaken in LB at 30 ^o^C or 37 ^o^C overnight. The overnight culture was brought back to exponential phase by diluting 70 μl culture into 4 ml fresh LB and shaking for 3h at 30 ^o^C. Frames and coverslips (Thermo scientific) were used to form an agar pad using 1% low-melting point agarose in PBS. Exponential phase cells were centrifuged at 6000 rpm for 1 min and resuspend in fresh LB. One microliter of mixed cells (v:v ratio = 1:1) was spotted onto the agar pad, allowed to dry, and then covered with a coverslip. The fluorescent labeled cells were imaged in each of two fluorescence channels (mTFP) and mKO) every ten minutes using a 63x oil immersion objective lens on an inverted wide-field fluorescent microscope (Zeiss AxioObserver.Z1). Acquired images were processed with customized Matlab scripts.

### Light Sheet Fluorescence Microscopy

Light sheet microscopy was performed on a home-built light sheet microscope based on the design of Keller et al (27) and described in (29, 56). In brief, the beams from either of two continuous-wave lasers (Coherent Sapphire, 448 nm and 561 nm) are rapidly scanned using a galvanometer mirror and demagnified to create a thin sheet of excitation light perpendicular to and at the focus of an imaging objective lens. The specimen is moved through this sheet in one-micron steps and fluorescence emission is captured to create a three-dimensional image. To image the entire larval zebrafish gut, four sub-regions are imaged and later manually registered and stitched. All exposure times were 30 ms and excitation laser power was set to 5mW measured at the laser output. A 5.5 Mpx sCMOS camera (Cooke Corporation) was used for all light sheet imaging, and a 40x 1.0NA objective lens (Zeiss). For time series imaging, scans occurred at 20-minute intervals for 12-17 hour durations.

### Sample Handling and Mounting for Imaging

Specimens were prepared for imaging as previously described in (29). Larval zebrafish were anesthetized using tricaine methanesulfonate at 120μg/ml, placed in melted 0.5% agarose gel at no more than 42 ^o^C, and pulled individually into glass capillaries. Each capillary was then mounted on a holder on a computer-controlled translation stage, and each fish was extruded in a plug of gel into a specimen chamber filled with sterile embryo medium and tricaine methanesulfonate. The fluid in the specimen chamber was maintained at 28 ^o^C. All time series experiments were performed overnight beginning in the evening.

### Imaging-Based Quantification of Bacterial Populations

*In vivo* gut bacterial populations were quantified from light sheet images using an analysis pipeline described in (29). In brief, bacterial aggregates and individual bacteria were separately identified. The number of bacteria per aggregate is estimated by dividing the total fluorescence intensity of the clump by the average intensity of individuals. Discrete individuals were detected using a wavelet-based spot detection algorithm, with autofluorescent host cells and other non-bacterial identified objects rejected using a support vector machine based classifier augmented with manual curation.

### Identification of Population Collapse Events

Collapses of bacterial populations were identified from light sheet microscopy time series images and visually confirmed as described in (30). Population collapses in *Aeromonas* were defined as a decrease in the total population of at least a factor of 10 in one time step (20 minutes), together with at least a factor of 2 decrease relative to the original population at the next time step, the latter to false positives from single bad datapoints.

### Intestinal Motility Measurements

Intestinal motility in larval zebrafish was imaged using Differential Interference Contrast Microscopy (DICM) as described in (31). Videos were recorded at 5 fps. A velocity vector field was determined from the image series using image velocimetry, and the amplitudes and frequencies of gut motions along the anterior-posterior (AP) axis were obtained using the analysis pipeline described in (45). In brief: The AP component of the vector field was averaged along the dorsal-ventral direction, resulting in a scalar motility measure at each position along the gut axis and at each time point. The frequency of gut contractions was calculated as the location of the first peak in the temporal autocorrelation of the motility. The amplitude was calculated as the square root of the spatially averaged power spectrum at the previously determined frequency, providing the magnitude of the periodic motion.

## Acknowledgements

We thank Rose Sockol and the University of Oregon Zebrafish Facility staff for fish husbandry, Sophie Sichel for preparation of germ-free zebrafish, and Karen Guillemin, Travis Wiles, and many members of the authors’ research groups for useful comments and conversations. Research reported in this publication was supported in part by the National Science Foundation (NSF) under awards 0922951 (to RP), 1427957 (to RP), and MCB-1149925 (to BKH); the M. J. Murdock Charitable Trust, and an award from the Kavli Microbiome Ideas Challenge, a project led by the American Society for Microbiology in partnership with the American Chemical Society and the American Physical Society and supported by The Kavli Foundation. This research was supported by the National Institutes of Health (NIH) as follows: by the National Institute of General Medical Sciences under award number P50GM098911 and by the National Institute of Child Health and Human Development under award P01HD22486, which provided support for the University of Oregon Zebrafish Facility. Funding for the work reported here was primarily from the Scialog Program sponsored jointly by Research Corporation for Science Advancement and the Gordon and Betty Moore Foundation through a grant to The University of Oregon, The Georgia Institute of Technology, and the Memorial Sloan Kettering Cancer Center by the Gordon and Betty Moore and Simons Foundations. The content is solely the responsibility of the authors and does not represent the official views of the NIH, NSF, or other funding agencies.

## Author Contributions

Designed research: SLL, JBX, BKH, RP

Performed research: SLL, JT, JY, RPB, DSS

Analyzed data: SLL, JY, JT, RPB, DSS, JBX, BKH, RP

Wrote paper: SLL, JT, JBX, BKH, RP

## References

1 Rajilić-Stojanović M, Heilig HGHJ, Tims S, Zoetendal EG, de Vos WM (2013) Long-term monitoring of the human intestinal microbiota composition. Environ Microbiol 15(4):1146–1159.

2 The Human Microbiome Project Consortium (2012) Structure, function and diversity of the healthy human microbiome. Nature 486(7402):207–214.

3 van der Waaij D, Berghuis-de Vries JM, Lekkerkerk Lekkerkerk-v null (1971) Colonization resistance of the digestive tract in conventional and antibiotic-treated mice. J Hyg (Lond) 69(3):405–411.

4 Buffie CG, et al. (2014) Precision microbiome reconstitution restores bile acid mediated resistance to Clostridium difficile. Nature 517(7533):nature13828.

5 Young VB (2017) The role of the microbiome in human health and disease: an introduction for clinicians. BMJ 356:j831.

6 Spees AM, Lopez CA, Kingsbury DD, Winter SE, Bäumler AJ (2013) Colonization Resistance: Battle of the Bugs or Ménage à Trois with the Host? PLOS Pathog 9(11):e1003730.

7 Barzilay EJ, et al. (2013) Cholera Surveillance during the Haiti Epidemic — The First 2 Years. N Engl J Med 368(7):599–609.

8 Silva AJ, Benitez JA (2016) Vibrio cholerae Biofilms and Cholera Pathogenesis. PLoS Negl Trop Dis 10(2):e0004330.

9 Laviad-Shitrit S, et al. (2017) Great cormorants (Phalacrocorax carbo) as potential vectors for the dispersal of Vibrio cholerae. Sci Rep 7. doi:10.1038/s41598-017-08434-8.

10 Halpern M, Izhaki I (2017) Fish as Hosts of Vibrio cholerae. Front Microbiol 8:282.

11 Almagro-Moreno S, Pruss K, Taylor RK (2015) Intestinal Colonization Dynamics of Vibrio cholerae. PLOS Pathog 11(5):e1004787.

12 Bartlett TM, et al. (2017) A Periplasmic Polymer Curves Vibrio cholerae and Promotes Pathogenesis. Cell 168(1-2):172–185.e15.

13 David LA, et al. (2015) Gut Microbial Succession Follows Acute Secretory Diarrhea in Humans. mBio 6(3):e00381–15.

14 Hsiao A, et al. (2014) Members of the human gut microbiota involved in recovery from Vibrio cholerae infection. Nature advance online publication. doi:10.1038/nature13738.

15 Russell AB, Peterson SB, Mougous JD (2014) Type VI secretion system effectors: poisons with a purpose. Nat Rev Microbiol 12(2):137–148.

16 Ho BT, Dong TG, Mekalanos JJ (2014) A View to a Kill: The Bacterial Type VI Secretion System. Cell Host Microbe 15(1):9–21.

17 Hachani A, Wood TE, Filloux A (2016) Type VI secretion and anti-host effectors. Curr Opin Microbiol 29:81–93.

18 Joshi A, et al. (2017) Rules of Engagement: The Type VI Secretion System in Vibrio cholerae. Trends Microbiol 25(4):267–279.

19 Bernardy EE, Turnsek MA, Wilson SK, Tarr CL, Hammer BK (2016) Diversity of Clinical and Environmental Isolates of Vibrio cholerae in Natural Transformation and Contact-Dependent Bacterial Killing Indicative of Type VI Secretion System Activity. Appl Environ Microbiol 82(9):2833–2842.

20 Sana TG, et al. (2016) Salmonella Typhimurium utilizes a T6SS-mediated antibacterial weapon to establish in the host gut. Proc Natl Acad Sci 113(34):E5044–E5051.

21 Anderson MC, Vonaesch P, Saffarian A, Marteyn BS, Sansonetti PJ (2017) Shigella sonnei Encodes a Functional T6SS Used for Interbacterial Competition and Niche Occupancy. Cell Host Microbe 21(6):769–776.e3.

22 Wexler AG, et al. (2016) Human symbionts inject and neutralize antibacterial toxins to persist in the gut. Proc Natl Acad Sci:201525637.

23 Chatzidaki-Livanis M, Geva-Zatorsky N, Comstock LE (2016) Bacteroides fragilis type VI secretion systems use novel effector and immunity proteins to antagonize human gut Bacteroidales species. Proc Natl Acad Sci 113(13):3627–3632.

24 Coyne MJ, Zitomersky NL, McGuire AM, Earl AM, Comstock LE (2014) Evidence of Extensive DNA Transfer between Bacteroidales Species within the Human Gut. mBio 5(3):e01305–14.

25 Russell AB, et al. (2014) A type VI secretion-related pathway in Bacteroidetes mediates interbacterial antagonism. Cell Host Microbe 16(2):227–236.

26 Milligan-Myhre K, et al. (2011) Study of host-microbe interactions in zebrafish. Methods Cell Biol 105:87–116.

27 Keller PJ, Schmidt AD, Wittbrodt J, Stelzer EHK (2008) Reconstruction of Zebrafish Early Embryonic Development by Scanned Light Sheet Microscopy. Science 322(5904):1065–1069.

28 Keller PJ (2013) Imaging Morphogenesis: Technological Advances and Biological Insights. Science 340(6137):1234168.

29 Jemielita M, et al. (2014) Spatial and Temporal Features of the Growth of a Bacterial Species Colonizing the Zebrafish Gut. mBio 5(6):e01751–14.

30 Wiles TJ, et al. (2016) Host Gut Motility Promotes Competitive Exclusion within a Model Intestinal Microbiota. PLOS Biol 14(7):e1002517.

31 Baker RP, Taormina MJ, Jemielita M, Parthasarathy R (2015) A combined light sheet fluorescence and differential interference contrast microscope for live imaging of multicellular specimens. J Microsc 258(2):105–112.

32 Fu Y, Waldor MK, Mekalanos JJ (2013) Tn-Seq Analysis of Vibrio cholerae Intestinal Colonization Reveals a Role for T6SS-Mediated Antibacterial Activity in the Host. Cell Host Microbe 14(6):652–663.

33 Ma AT, Mekalanos JJ (2010) In vivo actin cross-linking induced by Vibrio cholerae type VI secretion system is associated with intestinal inflammation. Proc Natl Acad Sci 107(9):4365–4370.

34 Lescak EA, Milligan-Myhre KC (2017) Teleosts as Model Organisms To Understand Host-Microbe Interactions. J Bacteriol 199(15):e00868–16.

35 Borgeaud S, Metzger LC, Scrignari T, Blokesch M (2015) The type VI secretion system of Vibrio cholerae fosters horizontal gene transfer. Science 347(6217):63–67.

36 Runft DL, et al. (2014) Zebrafish as a natural host model for Vibrio cholerae colonization and transmission. Appl Environ Microbiol 80(5):1710–1717.

37 Mitchell KC, Breen P, Britton S, Neely MN, Withey JH (2017) Quantifying Vibrio cholerae Enterotoxicity in a Zebrafish Infection Model. Appl Environ Microbiol 83(16):e00783–17.

38 Zheng J, Ho B, Mekalanos JJ (2011) Genetic Analysis of Anti-Amoebae and Anti-Bacterial Activities of the Type VI Secretion System in Vibrio cholerae. PLOS ONE 6(8):e23876.

39 Watve SS, Thomas J, Hammer BK (2015) CytR Is a Global Positive Regulator of Competence, Type VI Secretion, and Chitinases in Vibrio cholerae. PLOS ONE 10(9):e0138834.

40 Pukatzki S, et al. (2006) Identification of a conserved bacterial protein secretion system in Vibrio cholerae using the Dictyostelium host model system. Proc Natl Acad Sci 103(5):1528–1533.

41 Thelin KH, Taylor RK (1996) Toxin-coregulated pilus, but not mannose-sensitive hemagglutinin, is required for colonization by Vibrio cholerae O1 El Tor biotype and O139 strains. Infect Immun 64(7):2853–2856.

42 McNally L, et al. (2017) Killing by Type VI secretion drives genetic phase separation and correlates with increased cooperation. Nat Commun 8:ncomms14371.

43 Rolig AS, Parthasarathy R, Burns AR, Bohannan BJM, Guillemin K (2015) Individual Members of the Microbiota Disproportionately Modulate Host Innate Immune Responses. Cell Host Microbe 18(5):613–620.

44 Zac Stephens W, et al. (2015) The composition of the zebrafish intestinal microbial community varies across development. ISME J. doi:10.1038/ismej.2015.140.

45 Ganz J, et al. (2017) Image velocimetry and spectral analysis enable quantitative characterization of larval zebrafish gut motility. bioRxiv:169979.

46 Wiles TJ, et al. (2017) Modernized tools for streamlined genetic manipulation of wild and diverse symbiotic bacteria. bioRxiv:202861.

47 Foster KR, Schluter J, Coyte KZ, Rakoff-Nahoum S (2017) The evolution of the host microbiome as an ecosystem on a leash. Nature 548(7665):43–51.

48 Cremer J, et al. (2016) Effect of flow and peristaltic mixing on bacterial growth in a gut-like channel. Proc Natl Acad Sci 113(41):11414–11419.

49 Cremer J, Arnoldini M, Hwa T (2017) Effect of water flow and chemical environment on microbiota growth and composition in the human colon. Proc Natl Acad Sci 114(25):6438–6443.

50 Verster AJ, et al. (2017) The Landscape of Type VI Secretion across Human Gut Microbiomes Reveals Its Role in Community Composition. Cell Host Microbe 22(3):411–419.e4.

51 Bartlett DH, Azam F (2005) Microbiology. Chitin, cholera, and competence. Science 310(5755):1775–1777.

52 Pruzzo C, Vezzulli L, Colwell RR (2008) Global impact of Vibrio cholerae interactions with chitin. Environ Microbiol 10(6):1400–1410.

53 Tang WJ, Fernandez JG, Sohn JJ, Amemiya CT (2015) Chitin Is Endogenously Produced in Vertebrates. Curr Biol 25(7):897–900.

54 M. Westerfield (2007) The zebrafish book. A guide for the laboratory use of zebrafish (Danio rerio) (University of Oregon Press, Eugene, OR).

55 Choi K-H, et al. (2005) A Tn7-based broad-range bacterial cloning and expression system. Nat Methods 2(6):443–448.

56 Taormina MJ, et al. (2012) Investigating Bacterial-Animal Symbioses with Light Sheet Microscopy. Biol Bull 223(1):7–20.

